# Non-instructed Motor Skill Learning in Monkeys: Insights from Deep Reinforcement Learning Models

**DOI:** 10.1101/2023.12.04.569889

**Authors:** Laurene Carminatti, Lucio Condro, Alexa Riehle, Sonja Grün, Thomas Brochier, Emmanuel Daucé

**Affiliations:** Department of Informatics, Bioengineering, Robotics, System Engineering - University of Genova, Italy; Department of Robotics, Brain and Cognitive Science, Italian Institute of Technology (IIT), Genova, Italy; Institut des Neurosciences de la Timone (INT), Aix-Marseille Univ, Marseille, France; Institute of Neuroscience and Medicine and Institute for Advanced Simulation and JARA Institut Brain Structure-Function Relationships, Jülich Research Centre, Jülich, Germany; Ecole Centrale Méditerranée, Marseille, France

## Abstract

In the field of motor learning, few studies have addressed the case of non-instructed movement sequences learning, as they require long periods of training and data acquisition, and are complex to interpret. In contrast, such problems are readily addressed in machine learning, using artificial agents in simulated environments. To understand the mechanisms that drive the learning behavior of two macaque monkeys in a free-moving multi-target reaching task, we created two Reinforcement Learning (RL) models with different penalty criteria: “Time” reflecting the time spent to perfom a trial, and “Power” integrating the energy cost. The initial phase of the learning process is characterized by a rapid improvement in motor performance for both the 2 monkeys and the 2 models, with hand trajectories becoming shorter and smoother while the velocity gradually increases along trials and sessions. This improvement in motor performance with training is associated with a simplification in the trajectory of the movements performed to achieve the task goal. The monkeys and models show a convergent evolution towards an optimal circular motor path, almost exclusively in counter-clockwise direction, and a persistent inter-trial variability. All these elements contribute to interpreting monkeys learning in the terms of a progressive updating of action-selection patterns, following a classic value iteration scheme as in reinforcement learning. However, in contrast with our models, the monkeys also show a specific variability in the *choice* of the motor sequences to carry out across trials. This variability reflects a form of “path selection”, that is absent in the models. Furthermore, comparing models and behavioral data also reveal sub-optimality in the way monkeys manage the trade-off between optimizing movement duration (”Time”) and minimizing its metabolic cost (”Power”), with a tendency to overemphasize one criterion at the detriment of the other one. Overall, this study reveals the subtle interplay between cognitive factors, biomechanical constraints, task achievement and motor efficacy management in motor learning, and highlights the relevance of modeling approaches in revealing the respective contribution of the different elements at play.

**Author summary:** The way in which animals and humans learn new motor skills through free exploratory movements sequences solely governed by success or failure outcomes is not yet fully understood. Recent advances in machine learning techniques for continuous action spaces led us to construct a motor learning model investigate how animals progressively enhance the efficiency of their behaviors through numerous trials and errors. This study conducts a comprehensive comparison between deep learning models and experimental data from monkey behavior. Notably, we show that the progressive refinement of motor sequences, as they are observed in the animals, do not require the implementation of a complete model of their environment. Rather, it merely requires the capacity to anticipate both movement costs and final reward a few steps ahead in the future following a value iteration principle. Furthermore, the systematic deviations exhibited by the monkeys with respect to the computational model inform us on the presence of individual preferences in either minimizing the duration or the energy consumption, and also on the involvement of alternative “cognitive” strategies.

## 1 Introduction

### Motor skill acquisition in animals

The survival of animals in complex environmental conditions relies on their ability to hunt, forage for food, engage in fights, and crucially to continually learn and refine their skills to enhance both efficiency and competitiveness. As their bodies and environmental conditions change, the constant refinement of their motor abilities is an essential part of the learning process [1, 2]. Understanding the mechanisms underlying the acquisition and refinement of new motor skills has been an important focus of research in neuroscience.

Visuomotor reaching tasks have long served as a test-bed to study goal-directed motor control. They have provided a large body of evidence that under efficacy constraints, motor control should reflect the resolution of a certain speed/accuracy trade-off, that is finding the maximal speed at which a task should be completed through minimizing the error rate. The classical example of such a trade-off is observed in the famous Fitt’s task [3, 4], in which the maximum speed for alternating pointing movements between two symmetrical targets is regulated by the target size or the distance between the two targets. In humans and monkeys, visually-guided center-out reaching tasks have also been intensively studied with invasive and non-invasive recording techniques of brain activity to identify neural correlates of motor control and motor learning, under tightly controlled conditions [5–9].

In contrast with short term motor adaptation [10–13], motor skill learning takes place over longer time scales (from weeks to years), reflecting the slow acquisition of combination/coordination rules of the motor apparatus to control its many degrees of freedom [4, 14–17]. Complex skills generally require a specific combination of elementary motor commands [18–21], and involves visual/perceptual monitoring at different steps of the process [22–24], as it is the case for tool use in humans and primates, but also hunting or foraging in animals. The shaping of composite actions, made of many elementary movements, has conducted experimentalists to formulate the *chunking hypothesis*, which is the presence of a hierarchy in motor skill learning based on a composition of motor chunks [17, 20, 25, 26].

Importantly, the many degrees of freedom of the motor apparatus allow to achieve a singular task objective through different patterns of joint coordination [14, 27–29]. This can result in differences in strategies or performance between trials or individuals (intra and inter-individual variability) [30]. The adoption of a given solution by many individuals may also reflect its efficiency regarding a certain *criterion*. Classical motor learning models focus on optimizing kinematic criteria such as minimum jerk [31], variance control [32], energy cost [33], and achieving explicit goals through inverse models [27]. These principles of optimality readily combine with reinforcement learning models, enabling the interpretation of motor control as a trade-off between minimizing the costs of movement and achieving the task (e.g reaching a target) [34, 35].

Crucially, in the absence of supervision, the differences among trials for the same individual, may indicate that *different motor patterns*, matching the same criterion may be adapted in parallel, which implies that different combinations elementary movements chunks may allow to reach the same long term objective. s

### Free-moving task and Reinforcement Learning

In order to study more specifically the refinement of new motor skills, we need to analyze longer learning periods going from weeks to months, which is rarely the case in experimental settings. It is also necessary to develop free-moving and self-paced tasks in which subjects are allowed to explore a wide variety of motor strategies, with no explicit efficacy constraint.

Reinforcement learning (RL) algorithms were originally developed to resolve non-linear/non-inversible control problems in robotics [36]. Over the past decade, significant advancements have been made in the physical environment simulators [37], and have been coupled with novel methods that use auto-differentiation-based learning and gradient backpropagation within deep neural networks [38,39]. These have greatly improved the handling of continuous motor learning tasks [40–42], particularly self-paced motor tasks (e.g., learning of locomotor patterns [43]). Classically, RL agents learn to estimate the (cumulative) value of their actions from observing their final consequences, and retrospectively putting more weight on the actions for which to expect the highest total reward. The learning of long-term value relies on the principle of *Value iteration*, developed within the framework of dynamic programming [44].

### Proposed experiment and questions addressed

Leveraging on the combined knowledge from neuroscience, RL algorithms and the principles of optimality developed in classic motor learning models, we created this study to understand the mechanisms behind the acquisition of motor skills in monkeys performing a self-paced uninstructed reaching task.

The motor task used in this study was designed to explore the neural basis of complex motor behavior [45] and can be related to the implementation of a (simplified) traveling salesman problem. It requires to reach multiple targets in any order, from a central starting position in a 2D visuomotor space. Since the targets are always presented at the same location, the task facilitates motor exploration as well as planning and anticipation, beyond the classical constraints imposed by single target reach tasks [46].

Two macaque monkeys were trained to perform the task using positive reinforcement. Behavioral and neuronal data were recorded throughout the learning process [45, 46]. We then developed a reinforcement learning-based model to replicate the experiment, using a biomechanical simulator (that reproduces the conditions of the task), a parameterized controller, and a reward signal reflecting both the explicit reward feedback and the putative costs of task execution. Particularly, we tested 2 different cost signals: “Power” reflecting the energy spent to achieve the task, and “Time” corresponding to the total duration of a trial. This combination allows the interpretation of motor control as a trade-off between the achievement of the task (e.g reaching a target) and the minimization of the associated costs [34, 35]. In such a simulation, the RL agent’s performances can be compared with the ones obtained by the subjects. Moreover, key parameters like the reinforcement signal and costs can be freely asdjusted, providing a way to manipulate the causal relation between the guiding signal and the final control policies observed in experimental conditions.

In this study, we pursue two main objectives: (i) assessing the capacity of a deep reinforcement learning agent, specifically Soft Actor Critic (SAC) [42], to reproduce the monkey’s behavior, encompassing learning dynamics, motor pattern selection, and variability throughout training, and (ii) to elucidate the underlying factors influencing the monkey’s selection of specific motor patterns and refinement strategies.

We shed light on three fundamental issues in motor learning: (1) the relation between sequential motor patterns and the cumulative integration of reward; (2) the inclusion of kinematic efficacy criteria, such as the energetic/metabolic cost of the executed action, within the optimization process; and (3) the exploration of motor response variability across trials and individuals, and its significance in the optimization process.

## 2 Material and Methods

### 2.1 Monkey’s task

#### 2.1.1 Setup

This experiment involved two rhesus macaques (Macaca Mulatta): Monkey E, a 6 years old female weighting 7 kg and Monkey J, a 7 years old male of 9kg. They were trained to perform a task in which they had to reach and extinguish six simultaneously presented visual targets by displacing a cursor representing their hand position in the horizontal 2D space. Both monkeys were previously trained to perform visually guided reaching movements towards single target using the same setup. [45].

During the task, the monkeys were seated in a primate chair and held by a neck collar. Their right arm and forearm were supported by an articulated exoskeleton (Kinarm, BKin technologies, Ontario Canada) allowing for planar shoulder and elbow motion as well as independent monitoring of both joint’s motions as shown in Fig 1a at a 1kHz frequency. The non-working left arm was restrained in a semi-flexed position to avoid motor interference. This setup was coupled with a virtual-reality system composed of a horizontal semi-reflective mirror placed at chin height masking direct arm view, and a computer screen placed horizontally above the mirror and facing downward. The mirror could then reflect the screen’s display: the targets and a cursor (white dot) indicating at any time the position of the monkey’s hand in the workspace.

**Fig 1:**
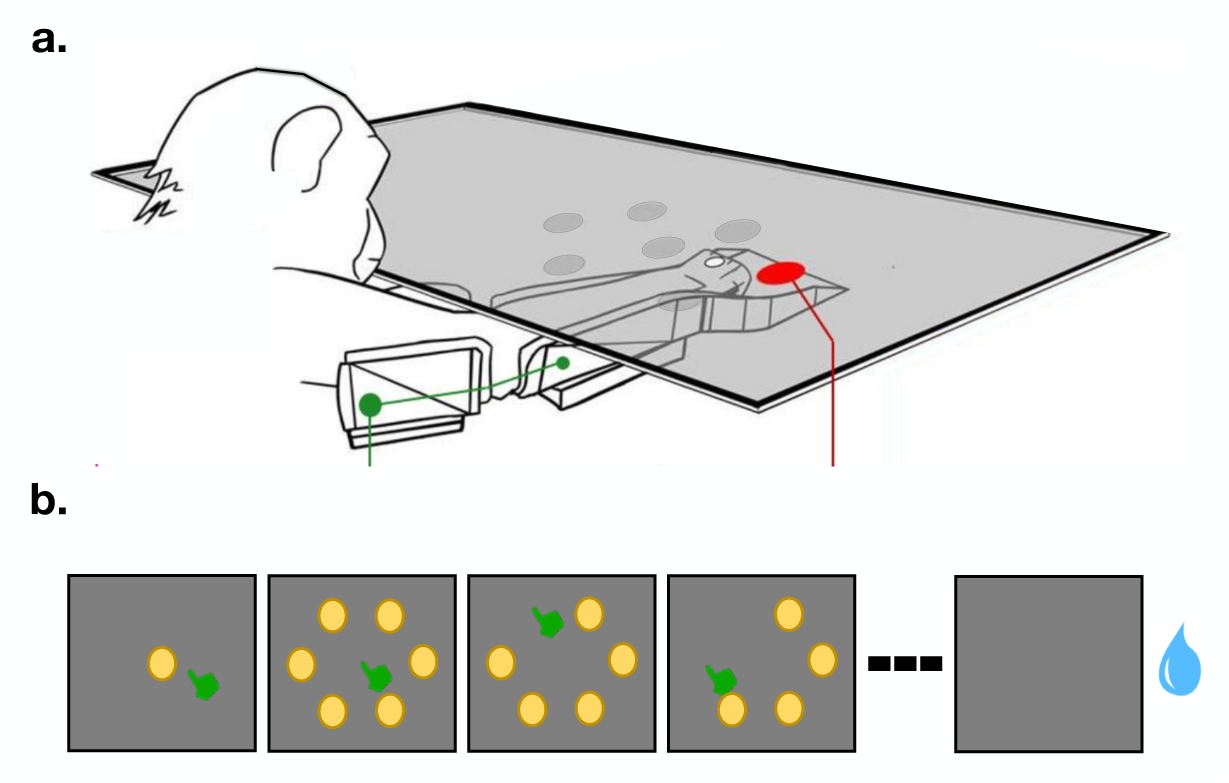
Self-paced multi-target reaching task. **a.** Experimental setup used to for the experimental task. Joint and arm kinematics were recorded at 1kHz (figure adapted from [45]) **b.** Visual display. Once the central target was reached, six peripheral targets appeared simultaneously on the screen. Each target was extinguished when reached by the hand feedback cursor. Once all the targets were successfully reached, the monkey was rewarded with a drop of water.

#### 2.1.2 Task and training

We use an original behavioral task in which the monkey had to perform controlled hand movements to reach several visual targets presented simultaneously. The hypothesis behind this task is that without any constraints on the order for target reach, the monkeys should develop integrated motor plans to optimize their hand trajectory along with the precision and speed of movement (Diamond et al., 2001).

The overall task consisted in reaching a set of 6 targets presented in the horizontal work-space. Each trial was initiated by the appearance of a central circular target (0.65 cm radius). Using the hand feedback cursor, the monkey had to reach this target and hold the position for 200 ms before the central target disappeared and six peripherals targets (0.65 cm radius) appeared simultaneously on the screen as shown in Fig. 1b. The six targets were displayed around the central one in a hexagonal configuration, and their location was adjusted to the maximal reach distance of each monkey. The target display remained the same throughout the experiment.

As soon as a target was reached, it disappeared from the screen. The interval between two consecutive reaches was limited to a maximum of 1500 ms, resulting in a total maximal duration of 9 seconds for the six targets. To promote swift movements, each target was considered “hit” once the hand reached a virtual circle with a radius of 1 cm centered on the target. Upon successful completion of reaching all targets, the monkeys were rewarded with water droplets 200 ms after the extinction of the final target. A new trial started 600 ms after the reward delivery. [45].

Pacing of task execution and order of target extinction were fully determined by the monkey itself. Although the monkey had ample time to reach each target within the time limit, reaching at faster velocity allowed to acquire the reward over more rapidly.

The spatial location of the peripheral targets differed slightly between the two monkeys as a function of their body size, being closer from each other for Monkey E than for Monkey J. The resulting “Setting 1” and “Setting 2” are described in the next section. During the entire learning period, Monkey E performed a total of 7,346 trials across 68 sessions lasting in total 23,872 s and Monkey J a total of 11,853 trials over 118 sessions for a total of 25,400 s.

### 2.2 Computational Model

We eveloped in Python a simulation platform that aimed at replicating the learning of the task by the two monkeys. The simulator reproduces the original 2-DOF Kinarm system (enhanced with actuators), as well as an adaptive controller. The controller consists of a multi-layer neural network, employing reinforcement learning principles to optimize the motor response.

#### Task Settings

To match the monkeys’ experimental conditions, we modeled two distinct settings (Settings 1 and 2) that employed the same target conditions and coordinate values as those used in the monkey experiments for monkey E and J, respectively. The target arrangement for both settings are depicted in Figure 2. The darker circles represent the visual targets and the lighter circles represent the non-visible active target area. This figure highlights the disparity in target distance for the two monkeys, with a larger spacing between targets in Setting 2 than in Setting 1.

**Fig 2:**
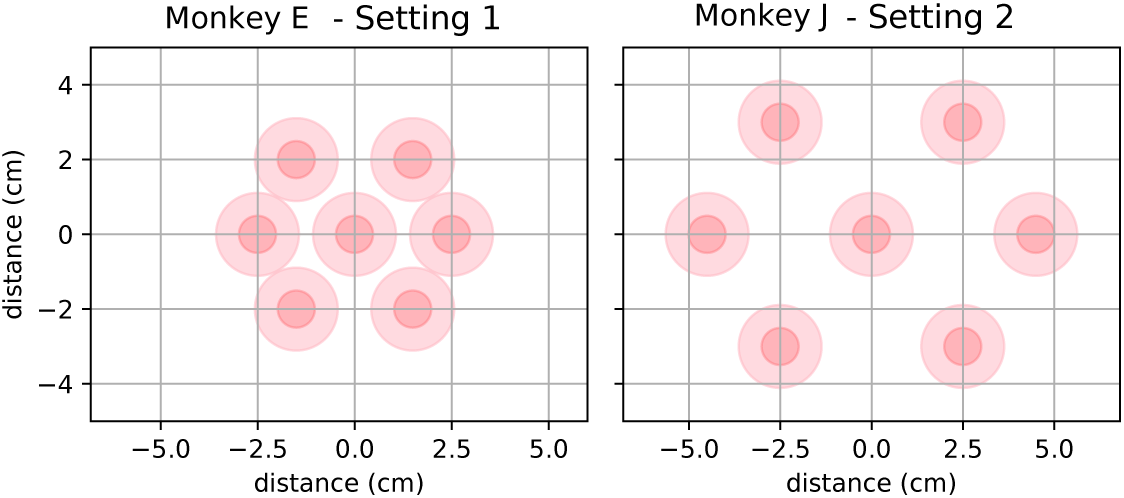
Representation of the targets arrangement of settings 1 and 2. The darker red circles depict the visible target as perceived by the monkey, while the lighter circles represent the (non-visible) active zone surrounding the target, which extinguishes the target when reached by the hand.

#### 2.2.1 Environment simulation

Our simulated environment conforms to the OpenAI Gym interface, largely used in reinforcement learning algorithms benchmarks [47]. In short, the simulated physical environment provides an interface that takes as input a motor command ***a***, runs the physical engine over a fixed period (here 500 ms), and returns the (updated) visual and proprioceptive information in the form of a state vector ***s***, a scalar reward *r* and a flag *d* that indicates whether the trial is achieved or not (Fig 3c). The physical engine is a custom first-order ODE integrator^1^, with a 30 ms temporal resolution.

**Fig 3:**
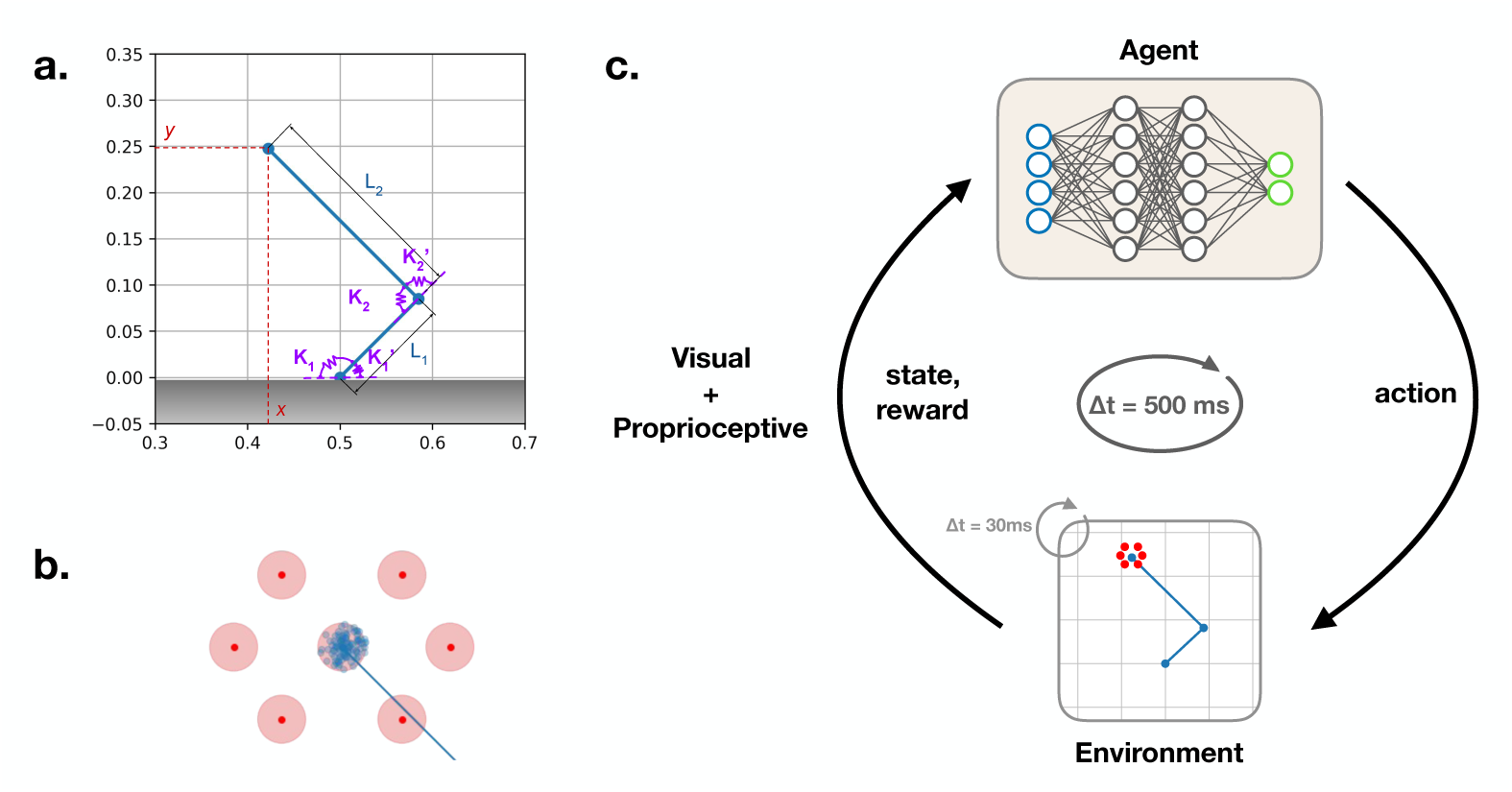
Simulation setup. **a.** Diagram of the 2-DOF mechanical model used in the simulations; (*x, y*) are the coordinates of the hand end position. **b.** The blue dots illustrate the distribution of the initial hand positions across trials. **c.** Schematic description of the closed-loop perception/action training setup, composed of an agent and an environment. The environment iterates the dynamics of the double-joint arm (with a 30 ms resolution), controls the disappearance of the 6 targets, and provides state observations and rewards at a 500 ms interval to the agent.

Our simulated environment uses a minimal mass-spring model [48], enabling the generation of a control in equilibrium point [31], directing the arm towards a unique position in space. The model as shown in Fig 3a is composed of two rigid segments: the upper arm and the forearm, with a total of two degrees of freedom: the elbow joint and the shoulder joint.

The dynamics of the two segments is controlled by a set of ordinary differential equations, that is inspired by the dynamics of a double inverted pendulum. The displacements taking place on a horizontal plane, the contribution of the gravity is null. The weight and length of the segments were adjusted to match that of a macaque’s arm. Each segment has two “springs” (anterior and posterior) for which the stiffness is modified to actuate a change in the angular position of the segment.

The state of the system is defined by the 2 joints angular positions *θ*_1_, and *θ*_2_, as well as their angular velocities 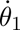 and 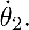 The length of the rigid segments are *l*_1_ and *l*_2_, and their mass is *m*_1_ and *m*_2_. The angular position of each joint *i* = {1, 2} is controlled by two antagonist forces: *F_i_*(*t*) and 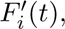 generated by two opposite springs whose stiffness are *K_i_*(*t*) and 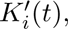 plus a damping factor *C*. When the stiffness are constant, the dynamics converges to an equilibrium point in the planar space. The final movement equations are thus the following:

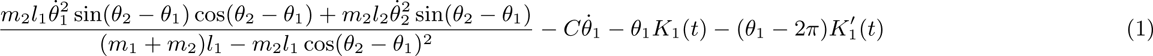

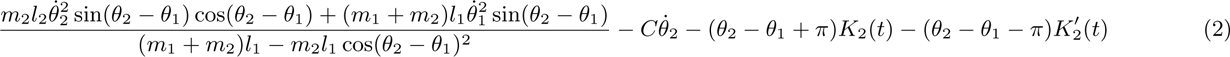

The practical values used in simulations are the following: *l*_1_=12 cm, *l*_2_=23 cm, *m*_1_=200 g, *m*_2_ = 500 g, *C*= 10 (damping factor).

To enhance the trial to trial variability as observed in animals, the hand initial position is normally distributed within a 1cm disc around the position of the central target, with a random initial angular velocity whose standard deviation is 0.1 rad *s*^−1^ (see Fig 3b).

##### Motor command

The upper arm and forearm are controlled by the agent through the stiffness of the springs 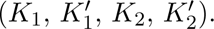 The motor command ***a*** = (*a*_1_*, a*_2_) defines a stiffness value for the four springs of the model, that is *K*_1_ = 12.5 + 2.5 *× a*_1_, 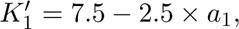 *K*_2_ = 5 + 5 *× a*_2_, and 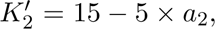 with (*a*_1_*, a*_2_) ∈ [*−*1, 1]^2^. Each command thus defines the value of two angular fixed points to which the arm should converge in the long run.

In our setting, the motor command is updated every 500 *ms*, reflecting a putative response time interval. Each execution step of our environment interface thus lasts 500 ms of the simulated physical time. The environment also records the disposition of the visual targets in the 2D physical space.

During an execution step, the targets disappear as soon as the extremity of the second segment (the “hand”) reaches the target within a radius *R* = 1 cm, at any time during the execution (the course of the hand is allowed to cross several targets during a single execution step). An example of the Kinarm hand surrounded by 6 targets is shown in Fig 3c.

##### Sensory input

At the end of an execution step, the environment interface returns a state vector ***s*** containing all visual, proprioceptive, and motor information.

In our implementation, the state of each target (lit or extinguished) is represented by a binary pair: the first element (”target on” cell) is set to 1 if the target is lit (and 0 otherwise), while the second element (”target off” cell) is set to 1 if the target is extinguished (and 0 otherwise). For the six targets, a total of 12 binary values thus represent the visual state.

For a total of *n* = 6 targets, the state vector is of size 2*n* + 4. It contains 2*n* binary values representing the state of the targets, plus the angular positions of the two joints, and a copy of the previous motor command (”efferent copy”).

##### Reinforcement signal

In the original task, monkeys are rewarded with a drop of water upon successfully extinguishing all six targets within the allocated time. We thus introduce in simulation a positive intrinsic reward granted at the end of a trial, when the controller effectively extinguishes all six targets. If the trial is completed within a time limit of 70*s*, the agent receives a reward equal to +10, else it receives 0. In the original task, the time-up limit was 9 s. In simulations, the time-up limit is extended to 70 s. Indeed, unlike the monkeys, the agent is perfectly ignorant of the task and the setup, while the monkeys were already trained in visuo-motor reaching task toward single targets using this apparatus. They were thus already aware that they had to touch the targets to receive a reward, while the model agents had to learn this association through random movements. Therefore, the longer time lime for the model was necessary in the ‘warm-up” part of the training, when the controller needed long durations of exploration to identify valid trajectories.

In addition to the final reward, we incorporated a penalty term (negative reward) that accounts for the costs associated with task execution. The goal of this study is to examine the impact of this penalty on motor behaviors acquired through learning. Consequently, we established two distinct learning conditions.

- In the “Power” condition, the negative reward equates the metabolic cost associated with arm control in space. By disregarding dissipative effects from damping, this energy cost can be quantified as the mechanical energy expended in controlling the arm. Specifically, the control of the springs stiffness constant induces a displacement that corresponds to a transfer of potential energy into kinetic energy. We describe this cost as the integrated kinetic energy of both segments of the arm over the time interval, defined as 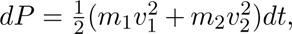 where *v*_1_ represents the velocity at the end of the first segment, *v*_2_ represents the velocity at the end of the second segment. This integrated cost is received at the end of each execution step and is scaled by a factor of 300 (so that the average penalty is ≃ 1 per second, depending on the velocity), i.e. 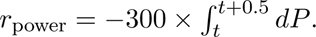
- In the “Time” condition, the algorithm incurs a penalty proportional to the elapsed time in seconds required to complete the task. The underlying hypothesis posits that the monkey strives to minimize task completion time, irrespective of the associated motor costs. In practice, this penalty is administered at regular intervals of 500 ms, resulting in a constant penalty of -0.5 applied after each execution step (so that the average penalty is exactly 1 per second).

#### 2.2.2 Motor control learning model

Reinforcement Learning algorithms train agents to find an optimal *policy*, that is a function that associates appropriate actions to the environmental *states*. The choice of the action is generally Markovian, that is independent of the past states. In other words, a RL agent does not require a model of the world but rather requires a correct sensory state (observation) *s* to provide the most appropriate action *a*. The goal of RL agents is to learn without explicit supervision, like the monkeys in our task.

At the heart of this learning method lies the fundamental principle of value iteration. Formally, value iteration considers a closed-loop control setup, where the current action *a* elicits an observation *s*^t^, which subsequently guides the selection of a new action *a*^t^, and so forth. Let *s* denote an observation, *a* the action taken, *r* the reward obtained, and *s*^t^ the state observed after the action. Furthermore, let *V* be a mapping from the state space to the set of real numbers. The value of the state *V* (*s*) can be recursively estimated using the following equation [36]:

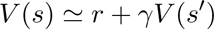

where *γ* is a discount factor in the case of open-ended interaction. The subsequent reward prediction error (TD-error) is the following:

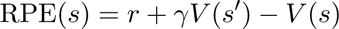

This allows the sensory guidance of action (online control) toward high values, but does not suppose to plan future actions in function of their expected outcomes (see also [49] for a discussion about model-based and model-free learning).

##### Soft actor critic (SAC)

Drawing upon behavioral observations that indicate a natural inclination of living organisms to continually explore their action space, the Soft Actor-Critic (SAC) algorithm is a RL algorithm that incorporates the objective of motor exploration into its optimization formula [42]. This objective takes the form of a maximum entropy criterion on the produced actions, combined with the long-term reward maximization objective. The entropy criterion on actions leads to stochastic policies, for which the algorithm employs various programming tricks to ensure gradient descent stability.

In practice, the controller is structured as two multi-layered perceptrons. The first network, referred to as the “actor”, takes the observation vector as input and computes the motor response. This network is coupled with a second neural network, known as the“critic”, which assigns a value to the actions produced by the actor. Both networks are trained simultaneously.

The inclusion of the entropy term in the optimization formulation relies on a temperature parameter, analogous to what can be found in stochastic optimization algorithms. This parameter, denoted as *β* (inverse temperature), is involved in the actor’s optimization formula as follows:

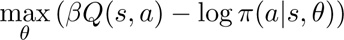

Here, *Q*(*s, a*) represents the action value function (the critic), and *π*(*a|s, θ*) represents the policy (or actor), viewed as a probability distribution over actions, and *θ* represents the policy parameters optimized by gradient descent (backpropagation). The inverse temperature parameter *β* controls the balance between exploration and exploitation, with higher values encouraging more exploitation (less exploration). By analogy with the behavioral observations showing a strong variability in the monkeys motor responses, the inverse temperature was kept constant (no annealing) during the full training and test procedures.

A replay buffer keeps track of the information (environment state - action - reward) needed to update the networks and learn. The training of the critic relies on a loss function that aims to maximize the accuracy of action value estimation using an entropy-biased Value iteration. It is noteworthy that, in the SAC procedure, the *Q*-value does not coincide with the Bellman optimum, but rather includes the action entropy term over the future pathway, i.e.

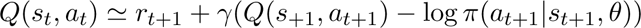

with *γ* being the discount factor. On the other hand, the training of the actor is based on the maximization of the value provided by the critic, augmented by the entropy term, as mentioned in the previous formula. This combination encourages exploration while balancing the exploitation of high-value actions.

By jointly training the actor and critic networks, the SAC algorithm leverages the actor’s ability to generate diverse actions and the critic’s ability to evaluate their quality. This interplay facilitates the learning process and enables the algorithm to discover effective motor sequences in continuous action spaces.

##### Training procedure

The SAC algorithm version employed in this study is based on the implementation provided by the OpenAI *Spinning Up in Deep RL* website [50]. The learning process is organized into trials called “episodes” (a sequence of movements up to extinguishing all 6 targets), wherein the controller interacts with the environment and receives sequences of observations and rewards. All these observations are recorded in a replay buffer and used as training data for the gradient backpropagation.

In order to provide empirical statistics, 10 learning agents were trained for each of the different task-condition combination, differing by the seed used for the random draw of the neural networks initial parameters. The two settings for the reaching task considered are presented in Fig 2 (Setting 1 and Setting 2). The two penalty conditions are the “Power” and “Time” penalties, as described earlier. Therefore, a total of 40 different agents were trained.

The training is structured into epochs, each encompassing 200 execution steps, equivalent to approximately 100 seconds of interaction with the environment. An epoch is not equivalent to a session, as the number of episodes (trials) in an epoch would depend of the average trial duration. Following each epoch, the controller undergoes a testing phase, during which the trajectory of the arm is recorded for one episode. The overall training spans 400 epochs, corresponding to a total of 40,000 simulated seconds (*∼* 11*h*) of interaction with the environment. This is to be compared with the 23,872 s recorded for Monkey E and 25,400 s recorded for Monkey J. We will therefore need to select windows of training that are comparable to monkeys.

The model’s hyperparameters were optimized to both achieve the learning of the task and provide hand trajectories approaching those observed in primates. Specifically, the optimizer used for the gradient backpropagation is Adam [51], the learning rate is equal to 10^−3^, and the hyperarameters are *β* = 5 (inverse temperature) and *γ* = 0.95 (time horizon discount factor).

Fig 4 presents the temporal evolution of the average cumulative reward (trial return) for the 10 agents of the 4 different conditions. The values account for both the 6-targets reaching bonus (+10) and the cost associated with the condition (energy or duration spent). The trend of the curve reflects the degree to which the controllers improve over time (heading toward value 7-8, with 10 being the absolute upper limit).

**Fig 4:**
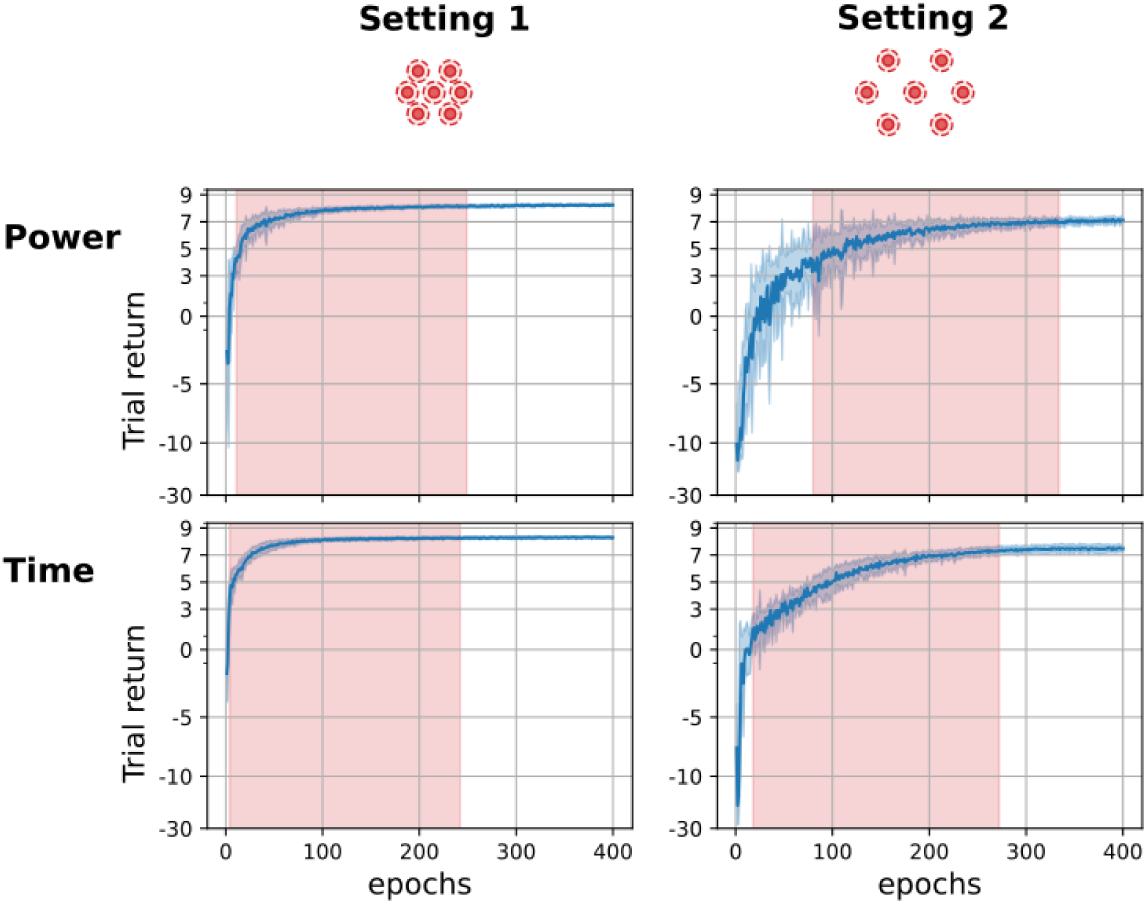
Learning curves. Mean evolution of the average cumulative reward (trial return) over the course of 400 epochs and across 10 agents, in the four conditions (Setting 1-Power, Setting 1-Time, Setting 2-Power and Setting 2-Time). The blue shaded regions indicate the variability, i.e. the episode return standard deviation among the agents. The red shaded regions represent the consolidation phase of the training used for comparison with the monkeys training data. A semi-logarithmic scale is used on the y-axis. Hyperparameters: *β* = 5, *γ* = 0.95.

For the purposes of analysis, the learning sessions were divided into three phases. The first phase is a warm-up phase characterized by rapid convergence, during which the agent learns to perform the task through random trial and error. The second phase is a consolidation phase (red shaded region), where it optimizes the efficiency of its motor response. Finally, there is a plateau phase, corresponding to the stabilization of the motor response.

The red shaded regions on the figure correspond to the consolidation periods that we considered for the comparison with the monkey’s learning data. These periods encompass the 240 epochs (24 000*s*) following the point at which the algorithm achieves the task in less than 6 movements on average (like monkeys at the start of their training). The initial learning period (“warm-up” period) is excluded from the comparison.

### 2.3 Data analysis

#### 2.3.1 Target reaching sequence

As for the monkeys, the order in which the targets are reached by the controller is not constrained. Consequently, the controller can choose freely the first target (one out of six), followed by subsequent target (one out of five), and so forth. We numbered the targets to record the sequences (order in which the targets are extinguished) and characterize and quantitatively assess the proportion of trajectories observed both in monkeys and simulations.

**Fig 5:**
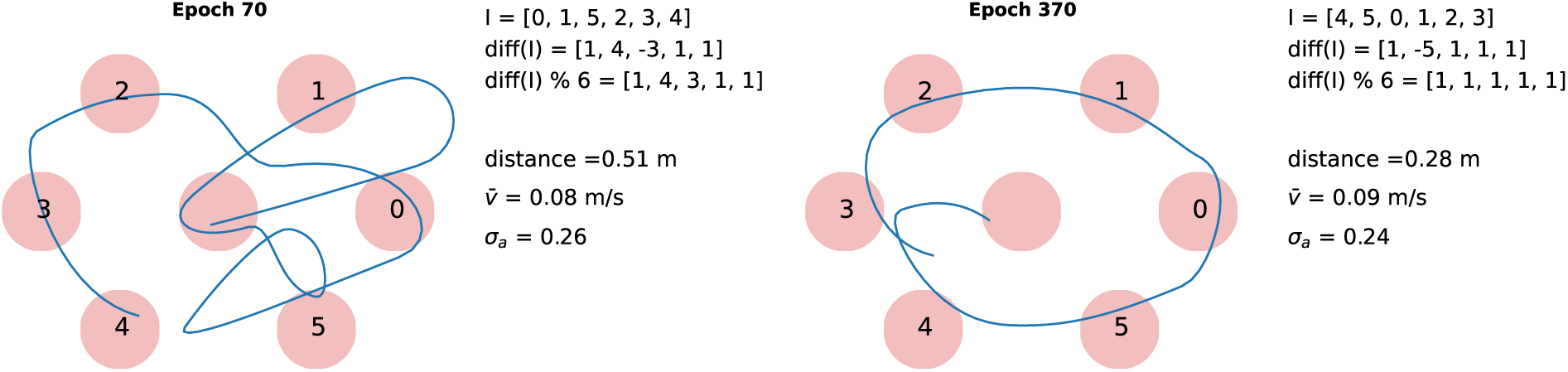
Circle vs. non-circles. The circularity of trajectories is measured by the difference in indices between consecutively visited targets within a single trial. If the sequence of these differences (modulo 6) is strictly equal to 1, then the trajectory corresponds to a circular visit in reverse clockwise order. Conversely, if the sequence of these differences (modulo 6) is equal to 5, then the trajectory corresponds to a circular visit in clockwise order.

Each target was assigned a number in counter-clockwise order, starting from the rightmost target. Subsequently, we analyze the sequence of these numbers as shown in 5: in the case of a counter-clockwise rotation, the difference between two consecutive numbers is 1 (modulo 6). Conversely, in the case of a clockwise rotation, the difference is 5 (modulo 6). Therefore, we identify any circular trajectory when the sequence of index differences (modulo 6) is either strictly equal to 1 or strictly equal to 5.

From this series of index differences, it is possible for each trial (or episode) to determine if a circular trajectory is employed to perform the task. For a series of trials, we can therefore estimate the proportion of circular strategies used in contrast to other possible motor sequences. However, we also want to estimate the number of quasi-circles, where the trajectory is visually circular but one of more targets have been barely missed. To obtain a precise assessment of attempts made towards circular trajectories, regardless of whether the targets are successfully reached, we introduce a supplementary metric termed as ‘circle attempt’. This metric analyzes the sequence in which the targets would have been reached had the diameter of the targets been a little larger. For Setting 1, the diameter is augmented by 0.2 cm, and in Setting 2, it is increased by 0.4 cm. These adjustments correspond to about 10% of the inter-target distance.

#### 2.3.2 Movement segmentation

To characterize the evolution of hand trajectories (both natural and artificial) and identify potential chunking happening in the learning process, we need to define what constitutes a movement.

Under the constraint imposed by the Kinarm exoskeleton, hand movements are confined solely to the two-dimensional (2D) plane. Therefore, a one-to-one correspondence exists between the articulatory positions and the spatial position of the hand. The analysis of hand trajectories relies on tracking the positional trajectory of the hand in both coordinates (*x, y*) over time, spanning from the central point to the final target. Kinematic variables, including velocity and acceleration, are derived from this trajectory and used for comprehensive motion analysis.

Among the most distinctive changes observed during the monkey’s learning process is a shift from a sequential ‘target-to-target’ reaching phase, where the monkey aims at each target in succession, to a more cohesive phase where it seamlessly engages with multiple targets within a single fluid motion. At first, the monkey aimed at one target after another, slowing down as it got close to each target and then speeding up again for the next one. Later on, the monkey started hitting multiple targets in a single smooth motion. Then, speed changes are much less pronounced, and the hand’s endpoint maintains a high velocity when passing through the target, while undergoing a deflection to change direction.

In the context of the artificial controller, the command results in a ballistic movement directed towards an endpoint. This command is updated every 500 ms. We hypothesize that complex integrated movements (such as the ones developed by the monkeys) require a precise control over directional changes by dynamically adjusting the endpoint. To distinguish such integrated movements, we segmented the hand trajectories using their velocity profile (see Fig 6). Our segmentation technique is founded on a dual threshold, estimated from the full trajectory. The lower threshold corresponds to the lower quartile of the velocity distribution, while the upper threshold aligns with the median velocity.

**Fig 6:**
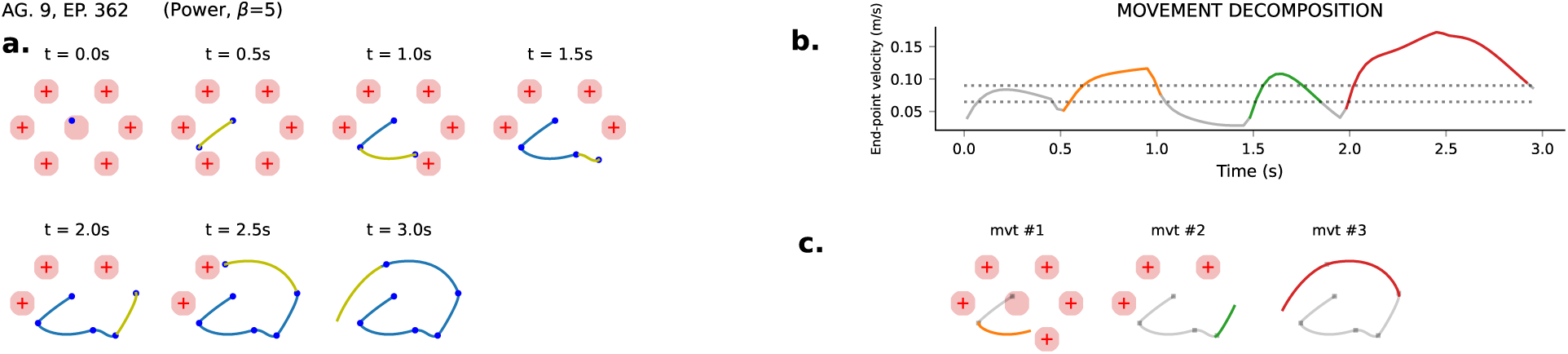
Movement segmentation. **a.** The hand movements are segmented based on the command update interval (every 500 ms). The trajectory consists of six sub-movements of 500 ms each, with progressive target extinction. **b.** Hand’s velocity profile across the entire trajectory. The lower threshold is set as the first quartile, while the upper threshold is defined as the median. In this case, three movements are identified, characterized by crossing the lower threshold, surpassing the upper threshold, and returning below the lower threshold. **c.** Movements segmentation reported on the hand trajectory, with a distinct color assigned to each movement, with corresponding targets extinctions.

In this way, an elementary movement is identified by a velocity sequence in which the velocity surpasses the lower threshold, rises above the upper threshold, and finally sinks back below the lower threshold. In short, a discrete movement is characterized by a clear initiation and termination, i.e. accelerating the hand, reaching a plateau and then decelerating significantly. The method is quite robust to local fluctuations and applies to a wide range of task and velocity conditions.

Based on this segmentation, each trial can be characterized by the number of movements needed to accomplish the task. An elevated number of movements (that is *n ≥* 6) should reflect a target-to-target reaching method, while a lesser number of movements (that is *n ≤* 5) reflects a more integrated multi-target reaching method. Integrated movements can be identified, distinguished by their longer duration compared to the update interval, as well as their curved trajectories that enable the attainment of multiple targets within a single movement (voir Fig 6c).

This method was used to differentiate between the initial phase and the consolidation phase, and select the learning window where the average number of movements is consistently below an average of six movements.

## 3 Results

In this section, the hand trajectories obtained for the models at different stages of learning are compared with those observed in monkeys. The dynamics and shapes of the trajectories are compared in relation to the two types of penalties considered in the simulation, namely the “Time” penalty (minimizing the execution duration) and the “Power” penalty (minimizing the total energy cost). To this end, we start with a qualitative evaluation of the shapes of the trajectories for both monkeys and models in each condition. We then use four different metrics, namely the number of movements, the covered distance, the trial duration and the average speed to quantify how the models depart or conform to the monkeys’ behavior.

### 3.1 Trajectories: qualitative aspect

Since the monkey and the model operate within the same 2D space, it is relatively straightforward to compare the evolution of trajectories between the two during the learning process. In particular, the control law can be characterized by its flow, which refers to the mapping that assigns a direction and movement speed to each point in space. Figure 7 provides a visual representation of this flow at various stages of learning, both in the case of the monkey and in simulation. This estimate is based on scattering the trajectories on 30 *ms* intervals, estimating the velocity in both directions, and representing the velocity with the direction of an arrow at each hand position, on 3000 observation points both in the monkeys and the models. By aligning 5 different flows at different stages of learning along a single row, we attain a comprehensive view of the behavioral changes attained throughout the learning process.

**Fig 7:**
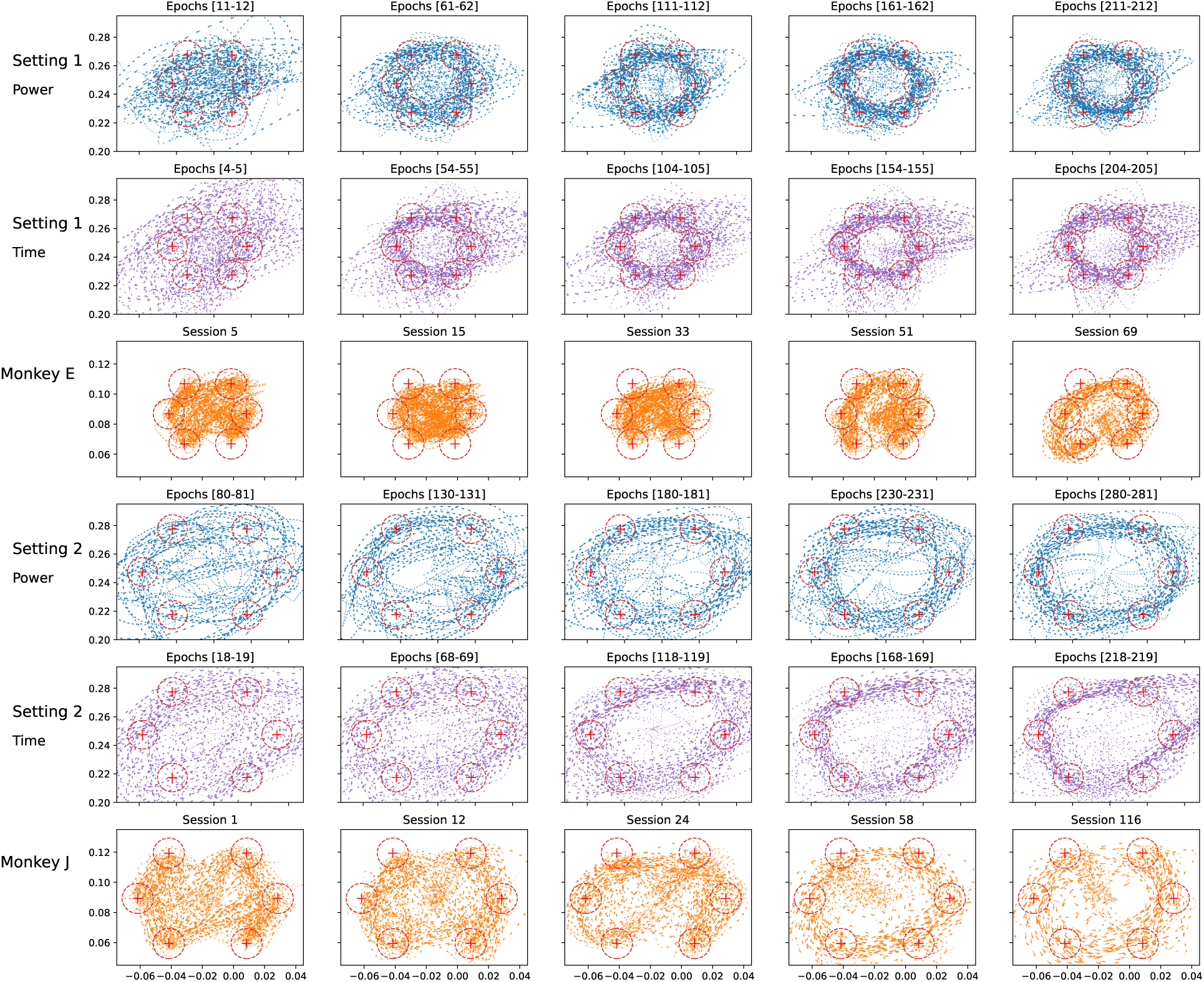
Superimposed endpoint trajectories recorded on 3000 time steps (picked over all the agents for the models on a given epoch interval) or 40 trials (picked in the same monkey session every 0.5 s), at different stages of learning. The sessions and epochs numbers increase from left to right, reflecting the motor control learning progress.

A similar trend is observed from left to right, from an initial tendency more disorderly, towards a more systematic and regular behavior with a circular aspect. This is observed as the trajectory passes through the visual targets in the same direction (counter-clockwise) for both the monkey and the model. This is notable as it reflects the presence of an optimization process at play, resulting in a convergence towards a relatively similar solution. In detail, the vast majority of the model trajectories are performed in an counter-clockwise fashion. Only one agent out of 10 converged to a clockwise response in the Setting 1/Power condition (not displayed in the figure), and is the only one showing a square-shaped trajectory.

A second important aspect is the maintenance of a significant across-trials variability. The behavior of the monkeys appears to be more noisy and less systematic compared to that of the models. A high diversity of behaviors is notably observed at later stages of learning, while the models exhibit considerably more reproducible behaviors. Noteworthy, Monkey J is more regular than Monkey E, probably due to the higher number of trials performed over the same total amount of time. Moreover, excursions beyond the task boundaries are observed within the models, especially under the “Time” condition, which is not seen in the monkeys.

Figure 8 provides illustrative examples of trajectories spanning the learning periods of both monkeys and models. These trajectories can be distinguished by their sequence (the order in which targets are reached) and by the overall trajectory shape. The comparison between the monkeys and the models is conducted here with a more fine-grained approach.

**Fig 8:**
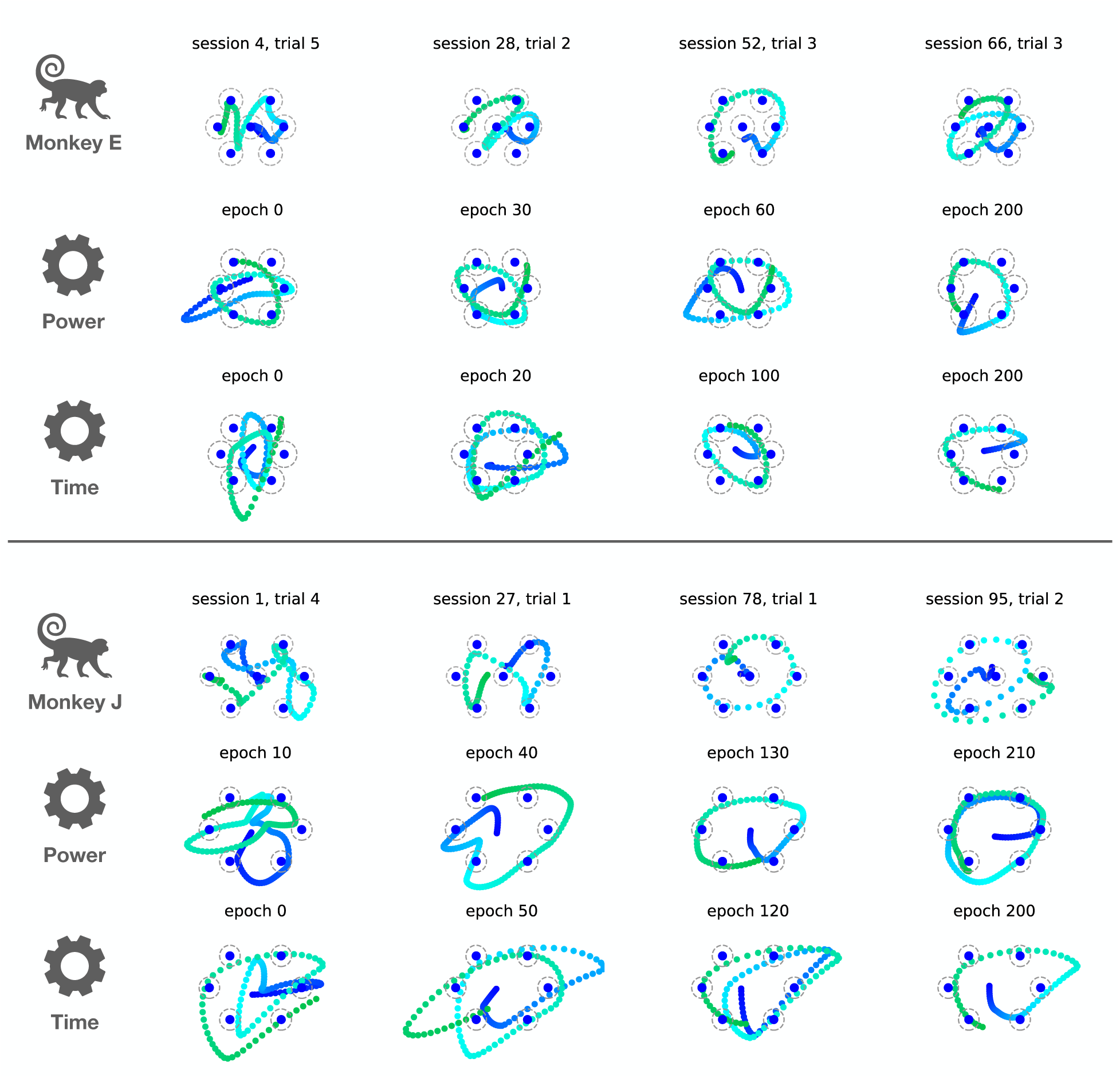
Example of trajectories of both monkeys and a chosen agent of each model condition, at different stages of learning. The sessions and epochs numbers increase from left to right, reflecting the motor control learning progress, for Setting 1/Monkey E and Setting 2/Monkey J.

Both monkeys start the experiment with a target-to-target reaching strategy. Over time, behaviors gradually evolve towards greater integration, wherein single movements are merged in a single trajectory targeting multiple consecutive targets. At the end of the experiment, both monkeys exhibit a more reproducible behavior with a majority of circles being performed in a smooth manner. The models exhibit a similar progression to that of the monkeys, transitioning from near-random behaviors to more integrated and circular actions.

Circular trajectories being the most prominent sequence in both models and monkeys reflects an inherent shortest path resolution. Interestingly, when a target is missed, the primarily intended sequence is often *repeated* with a slight inflexion, enabling the previously missed target to be reached, rather than performing a real-time online correction. This behavior, called a “double circle”, is illustrated in the last column of Fig. 8. It reflects a certain lack of flexibility, as if the full motor sequence was “automatized”. This strategy comes at a significant cost in terms of distance traveled, as the hand’s actual trajectory covers nearly twice the nominal distance.

Strikingly, models for both settings also exhibit this behavior rather than selecting a straight trajectory towards the remaining target. Overall, the models’ trajectories closely resemble those of their corresponding monkeys. However, the “Power” condition generally offers a closer match than the “Time” condition. Indeed, in the “Time” condition, the movement is significantly faster, especially in Setting 2, resulting in trajectories that are less centered around the targets. Additionally, all the trajectories presented here are performed in an counter-clockwise manner, confirming the observation made in Fig. 7.

Two important differences can however be observed between the models and the monkeys. The first one concerns the monkeys initial target-to-target strategy, reflecting pre-training, absent in the model. The second is that, unlike the models, the monkeys continue to produce a wide variety of trajectories that depart from circular movement even after intensive training. In the initial phase of the training, Monkey J predominantly performed trajectories following an infinity shape (*∞*), while Monkey E explored a larger number of non-circular trajectories. The second column of Fig. 8 displays the favorite alternative for each monkey. Models, on the other hand, tend to progressively converge toward a single circular strategy.

Overall, the analysis reveals a clear trend observed in both monkeys and models, to form and *stabilize movement sequences* that are then reproduced, with some variation, from trial to trial.

### 3.2 Trajectories: quantitative aspect

#### 3.2.1 Task variables

Circular sequences provide the most evident advantage by minimizing the overall distance traveled. This section focuses on determining the extent to which this sequence is selected, both in the monkey and in the simulations.

To analyze the observed trajectories, we characterized them by the sequence at which the targets were reached during a trial. We then reported in Fig. 9 the evolution in the proportion of true and attempt circles for each monkey and their associated models. The darker shades represent the proportion of real circles among the trials of a session, and lighter shades the cumulative amount of real and attempted circles (i.e. circular trajectories that missed at least one target).

**Fig 9:**
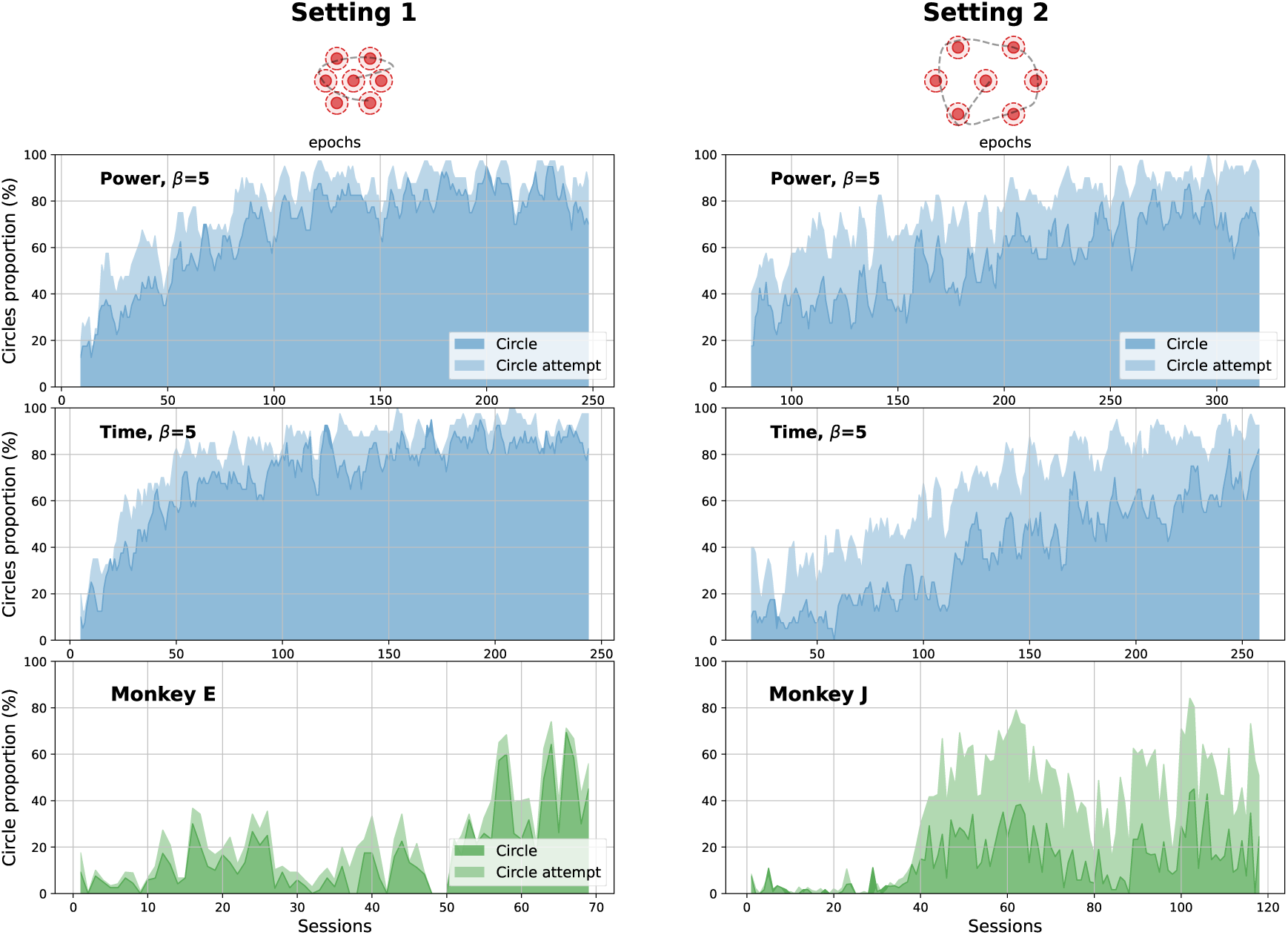
Circle proportion along sessions. Proportion (percentile) of circular hand trajectories observed across trials for each session (monkey) or for each corresponding epochs (model). Plain color: circles; shaded color: circle attempts. Left column: Setting 1/Monkey E. Right column: Setting 2/Monkey J.

As shown at the bottom of Fig. 9, both monkeys manifest a clear tendency to increase the number of circle trajectories during their learning process, but this trend is non-monotonic and characterized by abrupt transitions. Monkey E exhibits a relatively low and stable proportion of circles (between 0 and 30%) during the major part of the learning process, with a substantial increase around session 55, reaching a circle rate of approximately 60% in some sessions. Monkey J shows a clear biphasic evolution with an average of 5% true circles between sessions 0 and 32, followed by a sudden increase over 10 sessions, reaching an average of 20% from session 42 until the end of the experiment although with a notable dip around session 80. Moreover, Monkey J presents a large proportion of circle attempts, starting from 0% and reaching an average of 50% after session 40, being approximately twice the amount of true circles.

Unlike the monkeys, the models provide a consistent and regular increase in the proportion of circular trajectories. Moreover, all agents in all conditions converge toward almost exclusive circular trajectories. In Setting 1, both “Power” and “Time” conditions show an early increase in the proportion of true circles, although the “Time” condition converges a little faster to a majority of circles compared to the “Power” condition. In Setting 2, the evolution of the proportion of circles is rather linear in both conditions but with a lower starting value and a sharper increase in the “Time” condition.

Like the monkeys, the ratio of attempt circles is generally higher in Setting 2 models than in Setting 1, while the cumulative values at the end of the training (true + attempt circles) is the same between the settings, suggesting that the disposition of the targets in Setting 2 inherently leads to more missed circles for the same amount of total attempts. Table. 1 shows that Setting 2 agents in the “Time” condition also exhibit a significantly higher ratio of circle attempts than in the “Power” condition (t-statistic=-7.19, *p <* 0.001). In Setting 1 however, the difference is non-significant. Interestingly, for Monkey J, the amount of attempts circles is larger than that of real circles (ratio of 1.57), showing a closer similarity to Setting 2-Time.

**Table 1:**
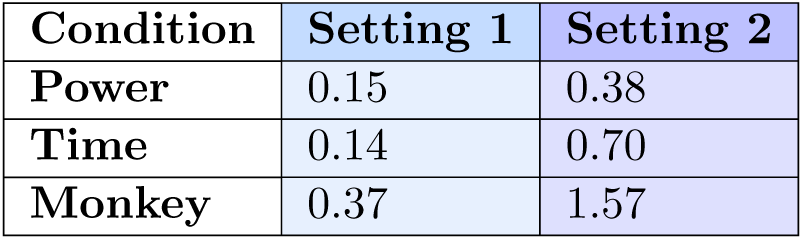
Table containing the ratio of missed circles to real circles in each condition and task setting. Numbers above 1 indicate a larger amount of attempts than real circles.

| Condition | Setting 1 | Setting 2 |
| --- | --- | --- |
| Power | 0.15 | 0.38 |
| Time | 0.14 | 0.70 |
| Monkey | 0.37 | 1.57 |

#### 3.2.2 Kinematic variables

Four kinematic variables were used to characterize and analyze the learning in monkeys and models:

1. the median *number of movements* necessary to reach all 6 targets (as explained in Section 2.3.2),
2. the average *distance* traveled by the hand during a trial,
3. the average *duration* of a trial and finally
4. the average *hand speed* during a trial.

Fig. 10 reflects the training of Monkey E and Setting 1 models. Monkey E is characterized by a relatively low hand velocity (less than 0.1 m/s) maintained during the full training. Its movement segmentation data (upper left panel) shows a tri-phasic evolution, with a fast decrease from 6 to 3-4 movements between sessions 1 and 20 (going from target-to-target to multi-target reaching), followed by a long plateau between sessions 20 and 60, and ended by an abrupt decrease from 3 to 1-2 moves between sessions 60 and 68 (Fig. 10a). Considering the hand traveling distance, once again a fast decrease (during the 5 first sessions) is followed by a steady value at around 0.2 m, never reaching the optimal shortest path distance, that is around 0.15 m for this task setting (dashed line) (Fig. 10b. This reflects a persistent variability in the movement as well as a certain proportion of missed targets, overall leading to longer pathways. Regarding the duration of the trials, the improvement follows a similar progression as the number of movements, implying that the increased chunking and smoothness of movement is the leading cause for duration reduction (Fig. 10c). The speed analysis also reveals an increased speed after session 60 concomitant with the decreased number of movements and duration (Fig. 10d).

**Fig 10:**
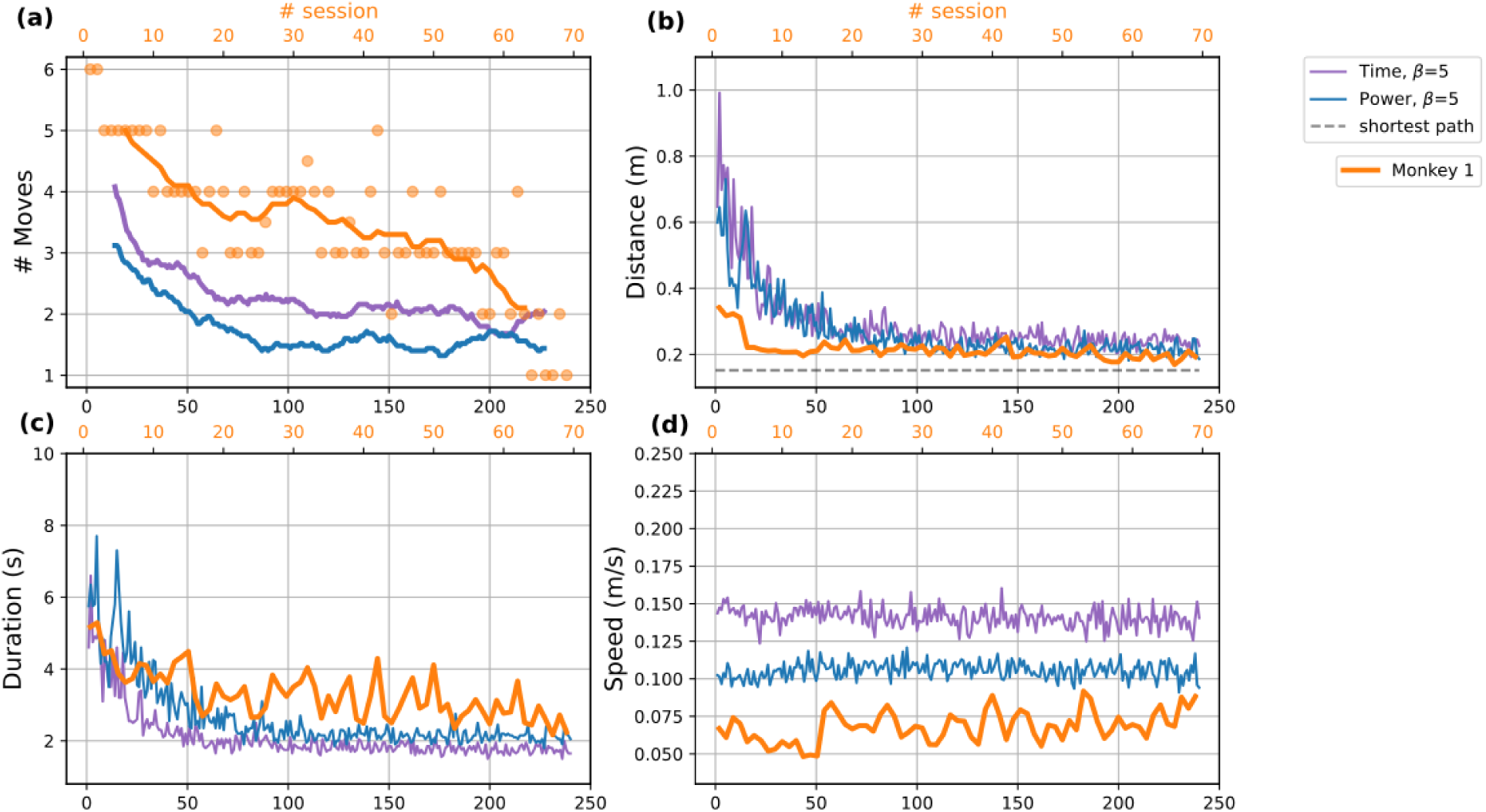
Evolution of characteristic kinematic variables across learning sessions, in Setting 1/Monkey E. This figure illustrates the progression of four variables (a) number of movements, (b) distance, (c) duration, and (d) speed throughout the training of models under both “Time” and “Power” conditions, aligned with Monkey E sessions.

Both conditions of the Setting 1 model exhibit an overall similar evolution profile across the four variables. However, their number of movements decreases more rapidly than that of the monkeys, with a final average value of 2 movements for the “Time” condition and of 1.5 for the Power condition. Learning mostly takes place within the first 50-60 epochs, accompanied by a reduction in the number of movements, distance traveled, and duration for both models. Interestingly, after this period, the distance traveled aligns with that of the monkey, reflecting a similar pathway optimization mechanism. The relatively low speed adopted by the monkey is not replicated in the models, which consequently perform the task faster than the monkey, with the “Time” condition being faster than the “Power” condition. In the case of both models and the monkey, learning appears to occur with little variation in average speed.

The learning process shows differences between the optimization of the monkeys and that of the models, but converges toward a very comparable solution. Interestingly, the final performance of the monkey (sessions 66-68), though slightly slower, hardly differs from that of the “Power” model. Overall Monkey E tends to behave closer to the “Power” condition, with a general reduction in the number of movements and duration along the training. It seems to be optimizing the precision of its movements, at relatively low speed, with a very low amount of missed circles.

Fig 11 provides us with the evolution of the same four variables along the learning process of Monkey J and Setting 2 models (in “Time” and “Power” conditions) for comparable learning windows. The first striking observation is the pronounced difference in dynamics between Monkey E and Monkey J. Indeed, the latter has a movement velocity that is two to three times faster. This leads to the fact that Monkey J is performing twice as many trials within a comparable duration, inducing a more extensive learning experience.

**Fig 11:**
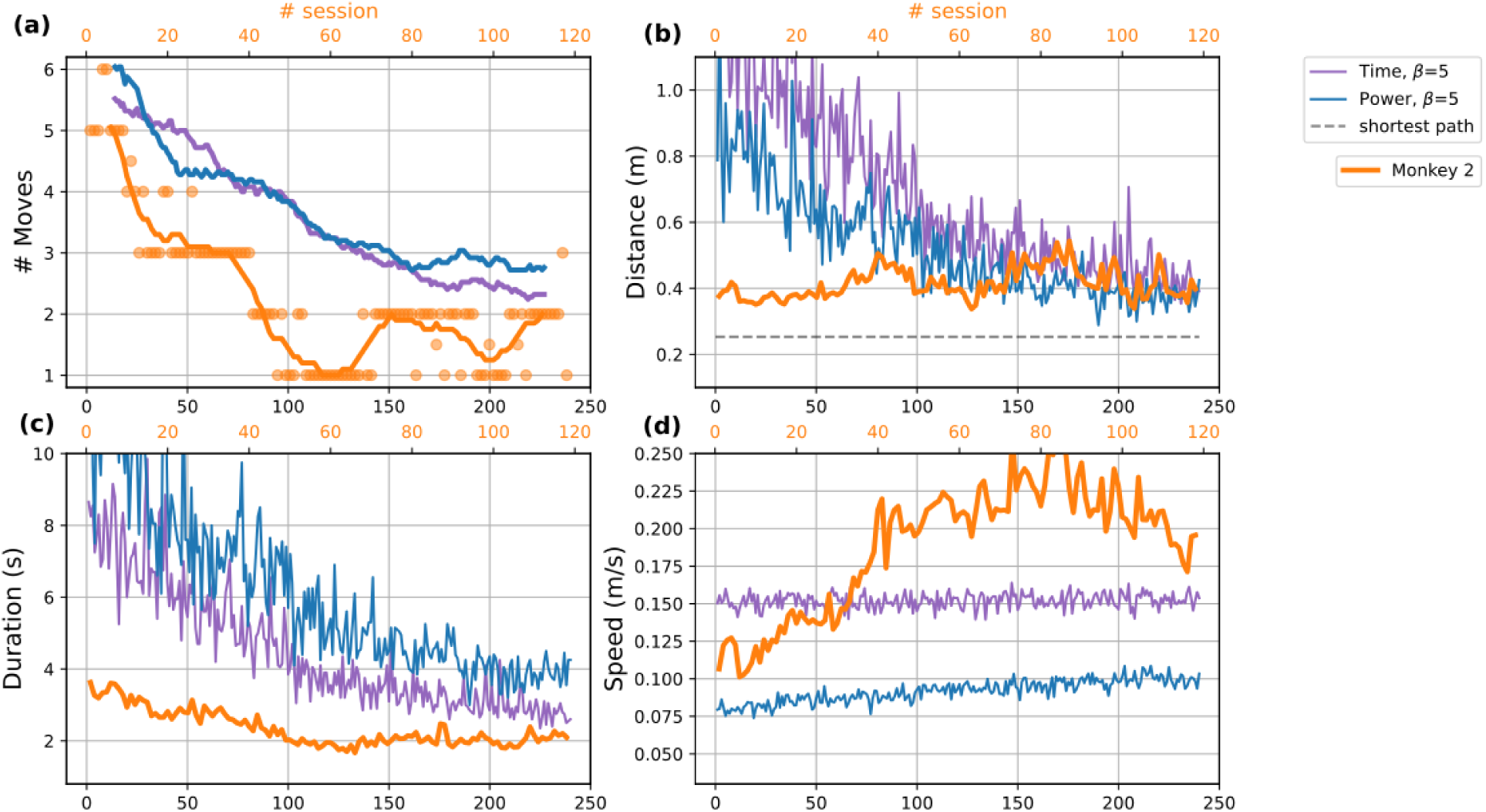
Evolution of characteristic kinematic variables across learning sessions, in Setting 2/Monkey J. This figure illustrates the progression of four variables (a) number of movements, (b) distance, (c) duration, and (d) speed throughout the training of models under both “Time” and “Power” conditions, aligned with Monkey J sessions.

In Monkey J, the evolution of the number of movements reveals interesting patterns characterized by the alternation of rapid decrease and plateaus, similar to Monkey E. There is a first rapid drop from 6 to 3 movements over the course of the first 20 sessions, followed by a plateau lasting until session 40. A second drop from 3 to 1-2 movements occurs over the 10 subsequent sessions followed by a variable plateau of a number of movements oscillating between 1 and 2 for the remaining sessions (see Fig. Fig 11a). As mentioned in the analysis of Setting 1, the presence of these plateaus can be interpreted as indicative of different “learning phases” for the monkey. Interestingly, and in contrast to Monkey E, the other variable undergoing the largest changes during the learning process is the speed, with a regular increase from session 1 to 80 sessions, followed by a decrease in the last sessions (see Fig. Fig 11d). The total distance covered by the hand during the trials shows minimal variations around 0.4 meters, with an increase to 0.5 meters coincidental with the speed peak (see Fig. Fig 11b. Overall the distance fails to reach the shortest path distance of approximately 0.25 meters. This can be explained by the high speed, leading to diminished precision and a larger amount of “double circles”. As for the trial duration, there is a continuous decrease during the first half of the learning process (up to session 60), followed by a stationary phase for the remainder of the learning period (see Fig. Fig 11c).

Regarding the simulations, once again, a relatively similar behavior is observed in both the “Time” and “Power” conditions. As opposed to Setting 1, the decrease in the number of movements in Setting 2 is more gradual and linear, converging towards an average value ranging between 2 and 3, which is higher than that of the monkey. On the other hand, the average distance traveled decreases gradually during the first 60 sessions and stabilizes around 0.4 meters in both conditions, reaching the same value as the monkeys, which is above the optimal shortest path distance. Nevertheless, some differences should be noted between “Power” and “Time” conditions. Particularly, the latter seems to host trajectories with an overall greater distance covered over a shorter time period (lower duration), accordingly with the notably higher speed of hand movement.

Linking these results with those presented in Fig 9, it becomes apparent that the dynamics of the trajectories of Monkey J are closer to those obtained with the “Time” condition. Indeed, in contrast to Monkey E, Monkey J seems to prefer a strategy based on optimizing time, with higher hand speed and lower precision. Nevertheless, with the exception of execution speed, both conditions yield a task resolution (movement coordination, circularity, maintenance of trial-to-trial variability) that closely resembles what is observed in the monkey.

Globally, these analyses show that each monkey learnt to perform their task while optimizing different aspects: 1) Monkey E favored precision and got close to the shortest path distance very quickly while only slightly improving the speed and duration throughout the learning period. Those results appeared closer to those of the “Power” condition which optimizes movements in order to avoid unnecessary energy loss. 2) Monkey J, however, favored speed at the expense of precision, with a large amount of missed circles, an increasing speed and a covered distance consistently higher than the shortest path’s distance. This seems to be in accordance with the “Time” agents that aim to optimize the duration of the task. Despite those tendencies, all monkeys and models converged towards a majority of circular trajectories, suggesting that both “Time” and “Power” can be used as optimization strategies to solve the task. However, although models still show substantial variability within and across agents, they tend to display greater consistency in their behavior compared to the monkeys. Notably, they show a more uniform evolution, leading to a more pronounced convergence towards circular patterns.

## 4 Discussion

Both monkeys and models hold interesting similarities in several aspects.

First, like the monkeys, the learning models asymptotically converge to a circular trajectory that approaches the mathematical optimum for this task (traveling salesman problem). The motor sequence gradually unfolds through trial and error, considering cumulative rewards via value iteration. This unanimous convergence, in both “Time” and “Power” conditions, indicates that both the duration and the energy cost adequately reflect the constraints of the task. Moreover, the progressive combination of short, discrete movements into larger ones (principle of binding motor “chunks” [52]), allowing to reach several targets at once, are reproduced in the models.

Second, monkeys and models also converge to almost exclusively counter-clockwise motions. In the models, counter-clockwise motions are largely dominant, since only one agent (out of 40) displayed a clockwise motion, which was also the only squared trajectory that was not smoothed out during learning. These results reflect the fact that, despite its simplicity, our models adequately adapt, like the monkeys, to the mechanical constraint posed by the Kinarm apparatus.

Third, both monkeys and models show a variability in their movement responses. For instance, when executing a circle with the hand, both monkeys and models perform it with some variations. More intriguingly, the overall progression toward more efficient movements may experience intermittent setbacks in the form of brief regression periods, marked by the execution of sub-optimal trajectories reminiscent of those observed at the initial stages of training, then followed by a return to the learnt behavior. This non-monotony is observed in the models, and even more pronounced in the monkeys.

Last, when the inter-trial variability leads to miss a target, both monkeys and models tend to opt for a “double-circle” rectification, which repeats with some inflection of the initial trajectory. This non-reactive behavior is characteristic of an “open-loop” motor response. This suggests that monkeys, like the models, seem to opt for a form of “open-loop” control, with limited contribution of the visual feedback. This is consistent with our initial assumption that motor learning may follow a form of “Value iteration”, that tends to develop systematic action sequence patterns over time.

However, beside the slight differences in variability between models and monkeys, a more important difference was observed in the number of hand motions developed during learning. Indeed, during learning, both monkeys alternate between different hand movement sequences to solve the task. Some may have their preference for a while, before the circular one being predominantly adopted. On the contrary, the models directly converge, in a progressive way, from random movements toward circles. All these results highlight an important difference between the monkeys and the models. This variability in monkeys might have mixed origins: 1) muscle tiredness, boredom and decreased interest for rewards are only possible in monkeys, and might lead to a desire of varying motion, and 2) our learning algorithm, despite owning an incentive to promote variability, tends to converge to a unique solution with growing accuracy, leading to more stereotyped movements towards the end of the training. While the models capture the main aspects of monkey’s learning, they do not entirely render the complexity of a macaque’s cognition, and lack internal body states to drive behavior. This has to be remembered when attempting to infer which out of “Time” or “Power” the monkeys try to optimize while learning their task.

Moreover, one of our initial questions, in relation to motor learning, was the role of movement cost in trajectory optimization. To adress this issue, we contrasted two penalty terms: “Time”, which reflects a form of “greedy” resolution, and “Power” that accounts for the energetic cost of movement. We observed that both penalty terms lead to solutions that are comparable from a kinematic perspective. However, taking into account the energetic cost results in trajectories that are slower, smoother, more homogeneous in speed, and with less deviation in the trajectory. Although the “Time” penalty yields a satisfactory response trajectory, those obtained in the “Power” condition are more realistic, suggesting that internal information regarding the energetic cost of these movements may contribute to motor response optimization in the monkeys. The energy constraint could play a much more prominent role when the task involves controlling a larger number of degrees of freedom with numerous redundancies.

While bearing in mind that the models presented here cannot fully depict the intricacy of the monkeys’ behavior, the Monkey E seemed to be generally acting closer to the “Power” agents, keeping a relatively low speed throughout the training and optimizing the distance as to be close to the shortest path. On the other hand, Monkey J seemed closer to the “Time” agents, with a strong increase in speed leading to optimized duration. However, their behavior at the end of their training is very intriguing. It seems as if each monkey exploited their own strategy (optimizing “Power” or “Time”) to the point of it becoming less efficient, to then incorporate the other constraint. This result is particularly interesting as it indicates that monkeys take into account both duration and energy consumption, but might start their learning by optimizing mainly one cost (at the detriment of the other), then observe general improvements, and later realize the limiting effect of optimizing only that cost. They are then capable of correcting their behavior and incorporate the other cost into their strategy to further improve their efficiency. This might explain why monkeys exhibit more varied behavior than models: they have the capability to selectively optimize one aspect at the expense of another (e.g., increasing speed to lower duration eventually decreases precision, which in turn increases the overall distance and energy spent) and later have the flexibility to modify their behavior to restore efficiency.

Finally, as the models operating in a “model-free” manner, they do not define an inverse model or an explicit goal. Monkeys seem to first employ a similar model-free, low-level learning of chunk-integration, leading to the progressive refinement of trajectories under efficiency constraints. Their relatively highly stereotyped trajectories also leans in the direction of model-free learning. However, they also show a second level of decision (learning strategy), involving the selection and evaluation of different motor sequences. This observation aligns with findings indicating a form of hierarchy in motor learning and decision-making framework [20, 53].

## 5 Conclusion

From a neuroscience perspective, our study sheds light on two aspects often treated separately in motor learning: the learning of motor sequences to produce coordinated behavior in time or space [18–21], and the consideration of the long-term reward in the plasticity process [54–56].

In particular, the main difference observed between our free-moving self-paced reaching task and visually guided sequential reaching movement is the presence of a significant variability in both the execution of motor trajectories (motor variability) and in the *choice* of hand motor sequences (hand trajectory variations). This mixed variability indicates the existence of multiple levels of control in the learning process.

- The most elementary level (combination of chunks) appears to be subject to a conventional “model-free” optimization process, solving a temporal-credit assignment problem at the motor command level, and partly considering the energetic cost of the command, resembling the principles implemented in most deep reinforcement learning algorithms for continuous control [40–42].
- The more integrated level of control (choice of a motor pattern among several possible patterns), which appears to be present in the monkeys only, would require the establishment of a hierarchical learning model (or meta-learning [57]), where different policies would compete. It is worth noting that the implementation of a meta-learning principle does not necessarily involve the presence of an inverse model and/or the definition of goals. The relatively stereotyped nature of motor responses observed in monkeys suggests a preference for model-free learning at multiple levels of control.

From the machine learning perspective, our study confirms the relevance of considering kinematic features (and energetic penalties) in refining motor learning. However, it also highlights the significantly higher flexibility of motor learning in animals compared to mere value iteration in the Bellman equation. Thus, to develop more powerful learning algorithms, it is necessary to take into account the observed variability both at the level of motor execution and at the level of the selection of motor sequences to perform.

Without the need to learn an inverse model, our observations rather suggest the implementation of flexible hierarchical mechanism that makes a choice among several motor programs to solve a task, and promotes adaptability to changing environmental conditions. These mechanisms should be able to account for the variability seen in motor learning, enabling the development of more versatile and robust algorithms for motor skill acquisition.

## Acknowledgments

The project was funded by the International Research Project “Vision for Action” (CNRS/FZ Juelich), the Deutsche Forschungsgemeinschaft (GR 1753/4-2, DE 2175/2-1) Priority Program (SPP 1665); the Helmholtz Association through the Helmholtz Portfolio Theme Supercomputing and Modeling for the Human Brain; the European Union’s Horizon 2020 Framework Programme for Research and Innovation (No. 720270, 785907). We are grateful to our colleagues, Frédéric Barthélémy, Shrabasti Jana, Alexander Kleinjohann for their scientific and technical support in carrying out this work. We would also like to thank Luc Renaud and Emilie Rapha for their technical support.

## Footnotes

1 adapted from https://matplotlib.org/2.0.2/examples/animation/double_pendulum_animated.html|

## Notes

### Competing Interest Statement

The authors have declared no competing interest.

